# Nucleoid openness profiling links bacterial genome structure to phenotype

**DOI:** 10.1101/2020.05.07.082990

**Authors:** Mahmoud M Al-Bassam, Oriane Moyne, Nate Chapin, Karsten Zengler

**Affiliations:** Department of Pediatrics, University of California, San Diego, La Jolla, California 92093-0760, USA; Department of Bioengineering, University of California, San Diego, La Jolla, CA 92093-0412, USA; Center for Microbiome Innovation, University of California, San Diego, La Jolla, CA 92093-0403, USA

**Keywords:** Nucleoid structure, Tn5 transposase, Nucleoid-associated proteins, Chromatin structure, H-NS, HiC, Transcription factor binding sites, POP-seq

## Abstract

Gene expression requires specific structural alternations in the nucleoid structure to enable the access of the transcription machinery into the genomic DNA. In prokaryotes, DNA binding proteins, including nucleoid-associated proteins (NAPs) and transcription factors (TFs), drive the change in structure and gene expression. Currently, studies of global NAP and TF binding are often hindered by the lack of appropriate epigenomic tools. Here, we present POP-seq, a method that provides *in vivo* genome-wide openness profiles of the bacterial nucleoid. We demonstrate that POP-seq can be used to map the global *in vivo* protein-DNA binding events. Our results highlight a negative correlation between genome openness, compaction and transcription, suggesting that regions that are not accessible to Tn5 transposase are either too compacted or occupied by RNA polymerase. Importantly, we also show that the least open regions are enriched in housekeeping genes, while the most open regions are significantly enriched in genes important for fast adaptation to changing environment. Finally, we demonstrated that the genome openness profile is growth condition specific. Together, those results suggest a model where one can distinguish two types of epigenetic control: one stable, long-term silencing of highly compacted regions, and a second, highly responsive regulation through the dynamic competition between NAPs and RNA polymerase binding. Overall, POP-seq captures structural changes in the prokaryotic chromatin and provides condition-specific maps of global protein-DNA binding events, thus linking overall transcriptional and epigenetic regulation directly to phenotype.

## INTRODUCTION

Genome organization is crucial to all life forms. In eukaryotes, histone oligomers organize the chromosomal DNA into nucleosomes of defined sizes, the building blocks of higher-order structures. By contrast, such well-defined structures are lacking in bacteria. Instead, a wide variety of poorly conserved nucleoid-associated proteins (NAPs) control the dynamic organization of the nucleoid and directly affect how genetic information is accessed, interpreted, and implemented^1,2,3,4^. Among the most widely studied NAPs is H-NS in *E. coli*^5^ and its functional analog, Rok in *B. subtilis*^6^, both known to have high affinity towards AT-rich regions^7,8^.

Omic technologies have revolutionized molecular biology by providing accurate measurements of molecular components, such as protein, RNA, and *cis*-acting elements. Yet, there currently exist few techniques for comprehensive identification and assessment of dynamic NAP binding and nucleoid organization *in vivo*. Implementation of HiC and similar methods have provided vital insights into the three-dimensional structure of the chromosome. However, HiC is limited by its technical and bioinformatic intricacy, the need for extreme sequencing depth, and the prerequisite for highly synchronous cell cultures, limiting its use to only a handful of selected bacteria^3,9,10,11^ and archaea^12^. As such, studies of the bacterial nucleoid at large remain challenging and therefore the effect of NAPs on global gene expression and nucleoid conformation remains poorly understood in the most abundant and diverse domain of life.

Here, we describe POP-seq (**<u>P</u>**rokaryotic chromatin **<u>O</u>**penness **<u>P</u>**rofiling **<u>seq</u>**uencing) as a method to interrogate changes in the openness of prokaryotic nucleoids associated with changes in the growth conditions and rapidly elucidate the state of bacterial genome organization. We present the results of POP-seq experiments carried out on the two major model bacteria, Gram-negative *E. coli* and Gram-positive *B. subtilis*. We compare our findings with previously published genomic studies to unravel the relationships between POP-seq measurements, NAP binding, DNA compaction, and gene expression. First, we show that POP-seq footprint signals are highly correlated with transcription factor binding sites (TFBS) and can potentially be used to identify novel TFBS in *E. coli* with both high reproducibility and high resolution. The POP-seq signals were also found highly correlated with both H-NS in *E. coli* and Rok in *B. subtilis*, suggesting that silencing of AT-rich genes by the binding of specialized NAPs is widespread within Gram-negative and Gram-positive bacteria. Through the integration of POP-seq with HiC and RNA-seq data, we unravel the role of the silencing NAPs in the epigenetic control of fast-response AT-rich genes. Our results suggest NAPs that bind AT-rich regions are fundamentally required for both Gram-negative and Gram-positive bacteria, despite the lack of amino acid sequence homology among these NAPs. Overall, POP-seq can provide an extensive map of protein-DNA binding events and genome-to-phenome associations in a fast and cost-effective manner.

## RESULTS

### POP-seq captures the open nucleoid of bacteria

To study the accessibility state of chromatin in bacteria, we employed a hyperactive Tn5 transposase. We performed our studies with *E. coli* as it is the most profoundly studied bacterium with copious omics data readily available in the public domain. The nucleoids of *E. coli* cells were first fixed with 1% formaldehyde to stabilize short-range DNA-protein interactions. The fixed cells were lysed and ions and small molecules, such as salts and sugars, were removed from the lysate by buffer exchange to eliminate any chance of modulating Tn5 activity. The lysate was diluted to 700 pg of DNA (equivalent to ∼1,500 *E. coli* cells), after which the fixed nucleoid was fragmented by Tn5 tagmentation and the resulting fragments were PCR-amplified to generate POP-seq sequencing libraries (**Fig. 1a**).

**Figure 1.**
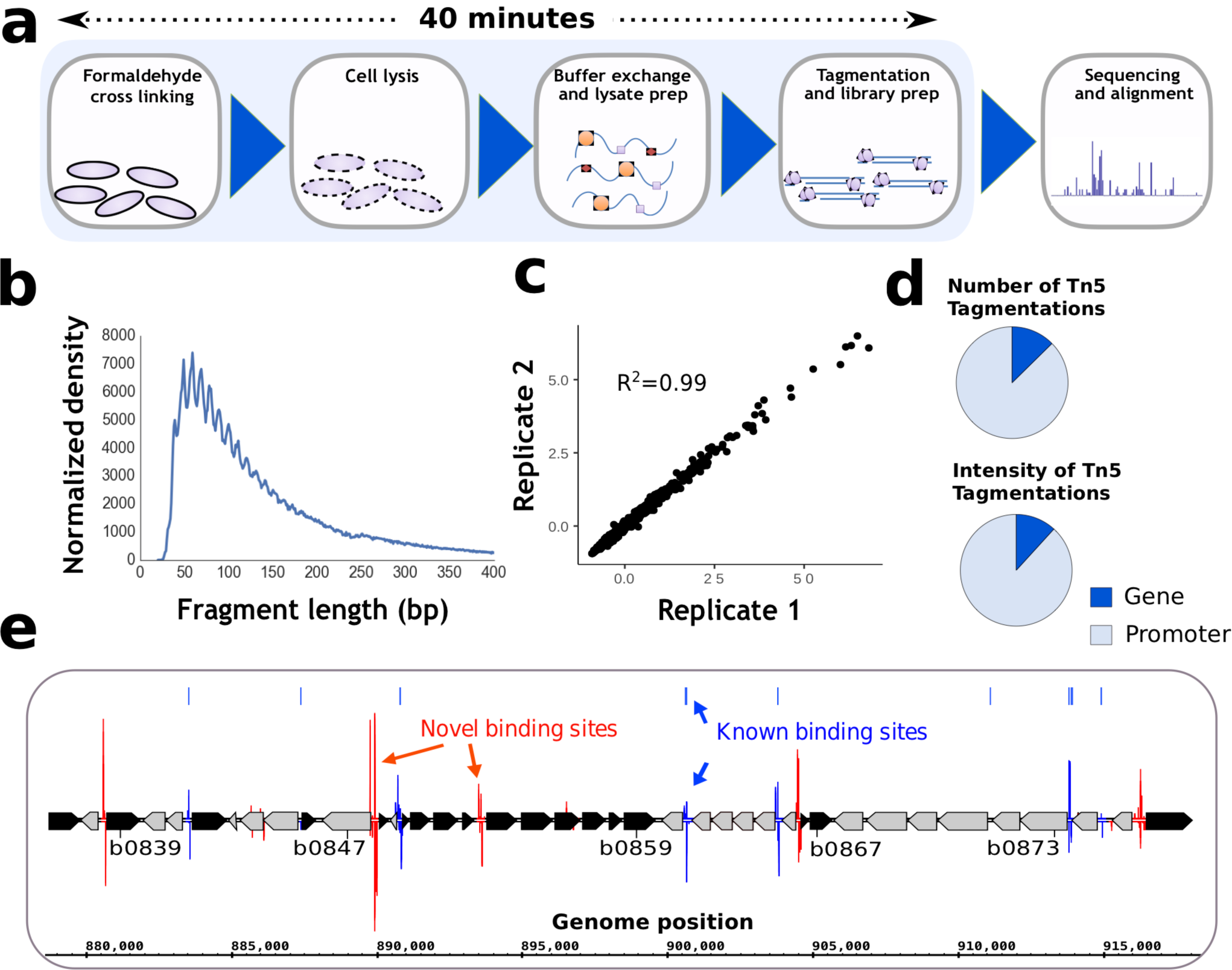
Overview of the POP-seq procedure and quality control checks of the data. **(a)** Summary of the major steps of the POP-seq method. The library generation takes ∼2 hours. **(b)** Distribution of the aligned fragment sizes. The subpeaks (spikes) are separated by ∼10 bp. **(c)** Correlation between two replicates of *E. coli* POP-seq. Each point represents the mean z-scored RPM coverage over a 5-kb window. The experiment is highly reproducible with R^2^=0.99 (p-value<2.2e-16). **(d)** Number of Tn5 tagmentation sites (top pie chart) and intensity of Tn5 tagmentation (bottom pie chart) found in the coding and the intergenic regions. The data is normalized by the total lengths of both regions. (**e**) Examples of POP-seq tagmentation sites. Most of the tagmentation events are present in intergenic regions. The blue spikes superimpose over experimentally verified TFBSs (small blue bars on top) from EcoCyc^17^. The red spikes could represent potentially novel TFBS

In eukaryotes, the fragment length distribution obtained from open chromatin has several notable characteristics that offer insight into the underlying chromatin structure. First, fragments generated from open chromatin were shorter than those produced from more compact regions. Second, the distribution of longer fragments fell into defined periods that reflect multiples of constant-sized nucleosomes. The POP-seq fragment length distribution ragned from ∼30 to ∼500 bp and was skewed to the right towards longer fragments (**Fig. 1b**), similar to eukaryotic fragment length profile^13^. However, no fragment periods similar to the ones found in eukaryotes^13^, were observed. The absence of this periodicity can be explained by the nature of the prokaryotic nucleoid, which lacks fixed sized nucleosomes. Instead, the prokaryotic nucleoid is organized and maintained by an array of NAPs^2,14^, leading to protected DNA fragments varying in size. Notably, the length distribution of both prokaryotes and eukaryotes had oscillations with a period of 10.5 bp, indicative of the helical pitch of DNA^13,15^.

POP-seq experiments performed on biological replicates were highly correlated (Pearsons’s R=0.99, p-value<2.2e^-16^, using average coverage over 5 kb windows), consistent with the concept that binding of NAPs and TFs is both highly organized and rigorously regulated (**Fig 1c**). The frequency of the Tn5 tagmentation events and the intensity of the resulting signals were greater at promoter sites compared to coding regions (**Fig. 1d**). It is well-established that DNA-binding proteins occlude Tn5 transposition at their sites of occupancy^13^, but the flanking regions are known to be hypersensitive to Tn5^13^ or DNase I^16^. Therefore, it is not surprising that strong POP-seq signals were found in intergenic promoter regions, particularly because these signals overlapped with known TF or NAP binding sites curated in EcoCyc^17^ (**Fig. 1e**). Interestingly, we found intense Tn5 signals at intergenic regions where no TFs or NAPs have been reported to bind, suggesting that POP-seq could detect novel regulatory binding sites in *E. coli* (**Fig. 1e**).

### POP-seq recapitulates previous NAP findings in bacteria

We found that AT-rich regions are hypersensitive to tagmentation by Tn5 **(Fig. 2a**). The broadly distributed POP-seq signal in these regions indicates that DNA binding events are occurring in a sequence-agnostic manner, highly reminiscent of the action of NAPs. In particular, H-NS is known for its strong tendency to bind to AT-rich regions^18^ (**Fig. 2b**). Genome-wide comparison of POP-seq signals with H-NS ChIP-seq signals revealed that the signals are highly correlated (Pearson’s correlation R=0.87, p-value<2.2 e^-16^ over 5-kb windows, **Fig. 2c**), suggesting that H-NS binding regions are hypersensitive to tagmentation by Tn5. Furthermore, POP-seq signals were enriched over broad-scale protein-rich domains (extensive protein occupancy domains, EPODs) previously identified by Vora et al., 2009 (magenta boxes in **Fig. 2d**). Overall, these findings imply that H-NS does not entirely occlude Tn5 transposase accessibility and that DNA flanking the H-NS binding sites is hypersensitive to tagmentation. This is consistent with *in vitro* experiments in which DNA bound by H-NS was resistant to DNase I digestion, while DNA immediately flanking the H-NS binding regions was hypersensitive to DNase I^19,20,21^.

**Figure 2.**
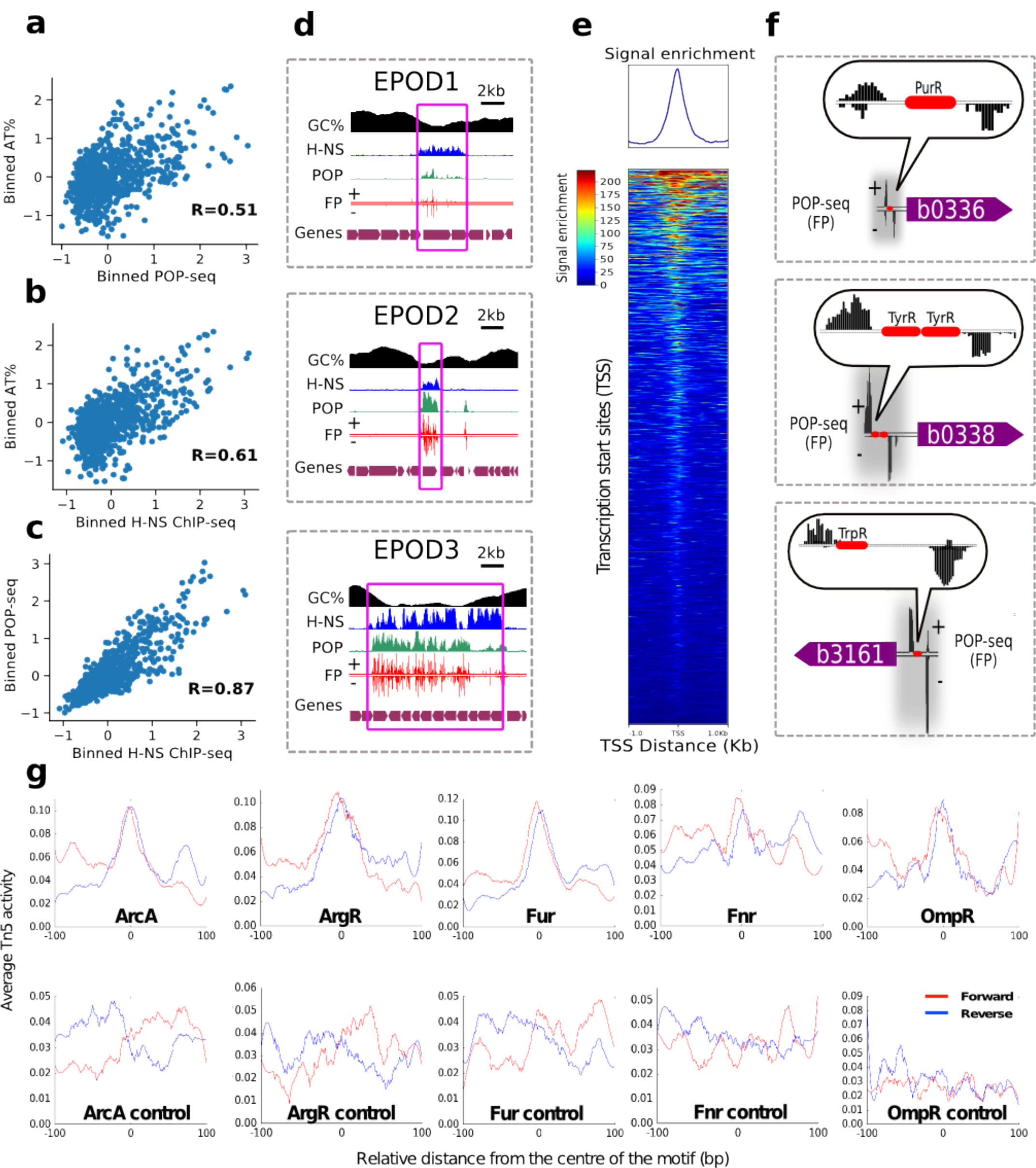
POP-seq recapitulates known TFs and NAPs binding regions. **(a)** Genome-wide correlation between POP-seq and AT% content datasets. **(b)** Genome-wide correlation between H-NS and AT%. **(c)** Genome-wide correlation between POP-seq and H-NS. The data in (a-c) were summed over 5 kb windows, z-score normalized and plotted. R represents Pearson’s correlation coefficient. **(d)** Agreement between POP-seq and protein occupancy domains (EPODs, Magenta boxes) reported by Vora et al. 2009. AT-rich regions are occupied by H-NS and are also accessible by Tn5 (POP-seq). The POP-seq footprint (FP) track is comprised of the 9-bp region between the two strand-transfer events catalyzed by each tagmenting transposase, which are by necessity occupied during tagmentation. It is therefore a more localized measure of accessibility to Tn5 than full-length reads. **(e)** Aggregated enrichment of the POP-seq signal over *E. coli* TSSs (top). The signal around each TSS (+/- 1 kb) was calculated and used to construct a heatmap in which the TSS-proximal regions are sorted by average signal strength (bottom). **(f)** Examples of PurR, TyrR, and TrpR TFBS from EcoCyc detected by POP-seq footprints. The binding sites are flanked by strong POP-seq signals on both positive and negative strands. **(g)** Cumulative footprinting signals for major TFs in *E. coli*. The Wellington algorithm^23^ was used to calculate the signals flanking the corresponding EcoCyc TFBSs. The control footprints over naked DNA shows no significant signals.

### POP-seq can identify transcription factor binding sites

The majority of gene regulation takes place at the transcriptional level. Transcription factors recognize specific sequences in *cis*-regulatory elements embedded in promoter sites and modulate transcription either positively or negatively. Due to the poor conservation of NAPs and TFs, the understanding of the regulatory network underlying gene expression represents a major challenge in bacteria. As such, we wondered if POP-seq signals can provide *in vivo* maps of DNA-binding events over the entire genome.

The median POP-seq signal over non-coding regions (including promoter sites) is higher than the median POP-seq signal over open reading frames (ORFs) by three-folds (Mann-Whitney p-value<2e^-16^). Further, we found that these signals are more prominent over transcription start sites (TSSs) (**Fig. 2e**), which is the center of transcriptional regulation. Our results imply that the nucleoid is more open at promoter sites, consistent with what is described in eukaryotes^22^, and that the signals could be originating from transcription factor-DNA binding events.

Thus, we examined the Tn5 tagmentation sites (POP-seq footprints), in which we only scored the 9 nucleotides covered by the Tn5 transposase for each aligned read. As a result, we generated two genome-wide alignment files for each experiment: one for the forward-strand sequencing reads and another for the reverse-strand sequencing reads. We tested if local depletions (footprints) in the two alignment files could be used to identify active transcription factor binding sites (TFBS) (**Fig. 2f**). We found that POP-seq signals are five-fold higher in the vicinity of putative TFBS curated in EcoCyc relative to the rest of the genome (Student’s t-test, p-value<2.2e^-16^). Further, we built a supervised model to predict whether each genome position is likely to contain a TFBS. As H-NS is a major contributor of the total POP-seq signal, the retained model took into account both POP-seq and H-NS Chip-seq signals, as well as the presence or absence of a gene at each genome position as predictors of EcoCyc-annotated TFBS (see Methods). With a sensitivity of 71% and a specificity of 84%, our model demonstrated that POP-seq is capable of efficiently highlighting genomic regions most likely to harbor a TFBS. Altogether, our results demonstrate that POP-seq can determine overall TF binding dynamics *in vivo*.

Next, we explored the POP-seq hypersensitive sites and tested if these can be used to determine TFBSs. We used the EcoCyc^17^ TFBS positions for each TF across the entire *E. coli* genome and explored the POP-seq signals that flanked the binding sites of each TF. Using the Wellington algorithm^23^, we found a sharp increase in the POP-seq signals flanking the center of many TFBSs tested (**Fig. 2g**), which diminished in positions distant from the TFBSs. The unfixed and naked DNA control showed no significant signals. These results demonstrate that POP-seq can reveal the positions of TFBSs with high accuracy.

### Highly accessible genes are readily adapted to growth conditions

In order to gain insights into the functional importance of genome openness, we performed RAST functional annotation of the *E. coli* genome, and tested which functional subsystems are associated with low or high POP-seq signals. Our results show that the genes with the lowest POP-seq signal are involved in house-keeping metabolic functions, such as ribosomal proteins, respiratory complex I, and electron transport complexes (**Supplementary Table 1**). On the contrary, genes with the highest POP-seq signal are involved in functions that require a fast adaptive response to changing growth conditions, such as alternative carbon source utilization pathways (e.g. xylose, D-ribose, D-allose, L-fucose, D-gluconate and ketogluconates), core-oligosaccharide biosynthesis, adherence and motility (especially related to fimbriae expression), CRISPRs, periplasmic acid stress response, toxin-antitoxin replicon stabilization systems, and general secretion pathways (**Supplementary Table 2**). Additionally, we observed that the 20 most Tn5-accessible subsystems present a significantly higher AT content and four-fold more H-NS binding than the 20 least-accessible ones (Mann-Whitney p-values<2.10^−16^). Together, these data demonstrate that genes that require a fast and reversible response to changing growth conditions are AT-rich, regulated by NAPs (H-NS in this case), and highly accessible to Tn5.

To validate the influence of bacterial growth condition on genome openness dynamics, we conducted POP-seq on *E. coli* grown in minimal medium (MM) containing ribose, xylose, or glucose. Principal component analysis (PCA) indicated that bacteria grown in MM+glucose or Luria-Bertani (LB) present similar POP-seq profiles, while remarkable differences were seen in MM+ribose or MM+xylose media (**Fig. 3a**). A global overview of the POP-seq signal measured in MM+glucose, ribose or xylose over the *E. coli* genome confirms the major differences between profiles, especially when comparing those obtained in presence (red lines **Fig. 3b**) or absence (green and blue lines **Fig. 3b**) of glucose as the main carbon source. These results highlight that POP-seq signals are directly linked to structural changes in the bacterial nucleoid and occur in direct response to changes in environmental conditions.

**Figure 3.**
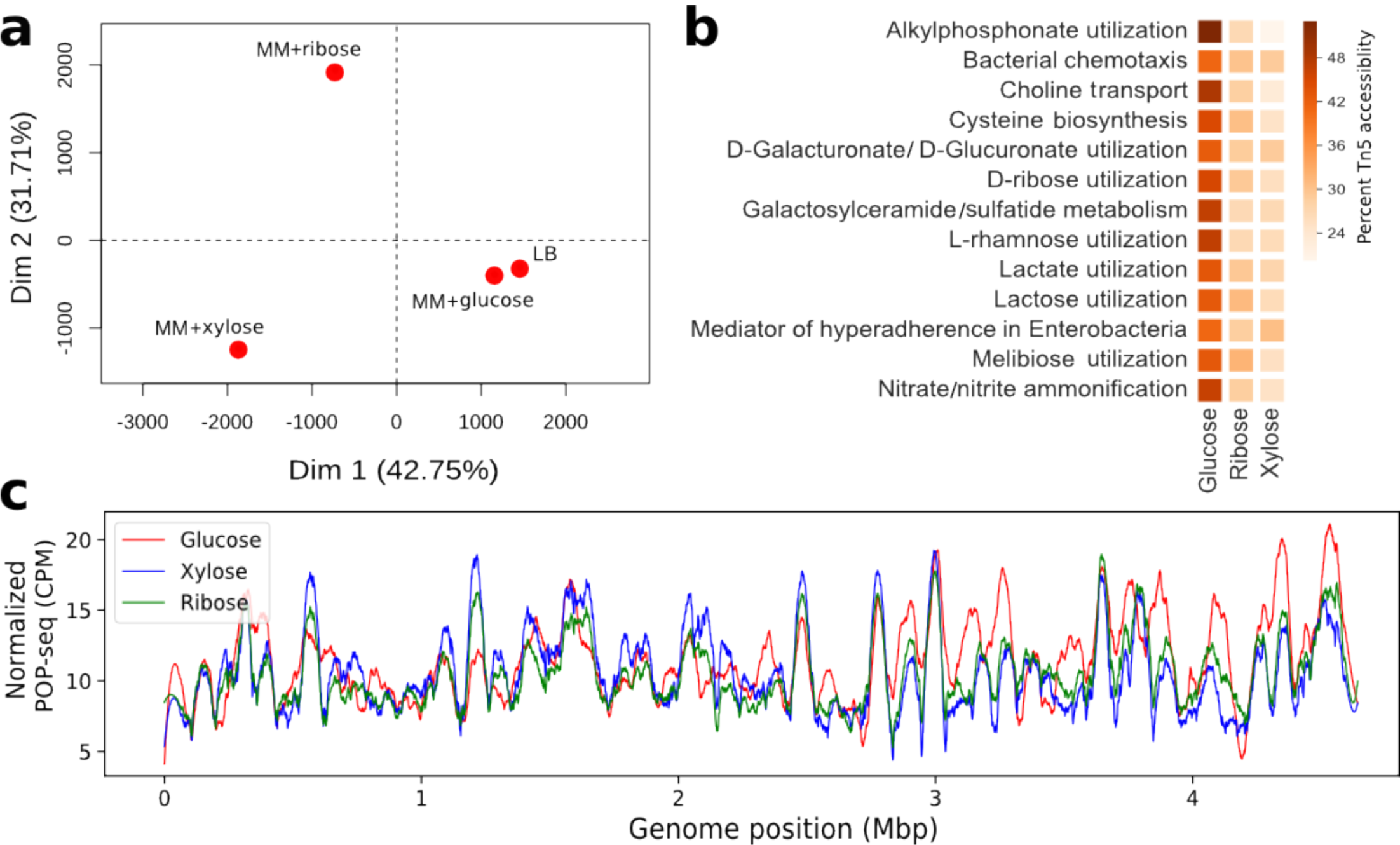
Differential POP-seq profiles of *E. coli* after growth in various culture media. **(a)** Principal Component Analysis of the mean POP-seq signal at every *E. coli* genome position. The overall POP signals are highly similar in MM+glucose or LB, medium while alternative carbon sources (ribose and xylose) induce major modifications. **(b)** Genome-wide POP-seq signal profiles of *E. coli* grown in MM with glucose, xylose, and ribose. The signals were smoothed using 100 kb windows. Glucose signals are significantly higher downstream of the 3 Mbp position **(c)** Heatmap showing all the RAST subsystems with a significantly lowered average POP-seq signal in both MM+ribose and MM+xylose compared to MM+glucose (DESeq analysis). The POP-seq values are normalized as percent values for each subsystem. Most of those subsystems are related to alternative sugar/carbon sources utilization pathways.

Moreover, comparative analysis allowed the identification of 13 RAST subsystems showing a significantly decreased POP-seq signal in both MM+xylose and ribose compared to glucose (DESeq2 p-values, FDR adjusted<0.05, **Fig. 3c**). Most of those subsystems are related to alternative sugar utilization pathways (**Fig. 3c**), suggesting a removal of NAPs transcription inhibition to adapt to the absence of glucose. These results show that POP-seq signals are directly linked to structural changes in the bacterial nucleoid, and occur in direct response to changes in environmental conditions. Therefore, POP-seq can link structural modifications of the nucleoid with function.

### The nucleoid openness is constrained by DNA compaction or transcription

Inspection of high- and low-accessibility regions shows that AT-rich genes are generally silenced by H-NS and have high Tn5 accessibility, while highly transcribed genes are far less accessible **(Fig. 4a)**, as elongating RNA polymerase occludes H-NS (and Tn5) accessibility^1^. Therefore, it is possible that high transcription or highly compacted genomic regions negatively affect genome accessibility measured by POP-seq. To assess this possibility, we evaluated the global relationship between DNA compaction^11^, RNA expression^24^, and Tn5 accessibility (POP-seq). To enable direct comparison, the 3-dimensional matrix of DNA-DNA contacts from a HiC experiment^11^ was flattened into a 1-dimensional array. All datasets were Z-score normalized and the signals were lightly smoothed and binned into 928 equal regions (5 kb each, **Supplementary figure 1**). We observed a significant negative correlation between POP-seq and HiC signals (Pearson’s R=-0.44, p-value<0.01), showing that highly compacted regions are not likely to be accessible to Tn5.

**Figure 4.**
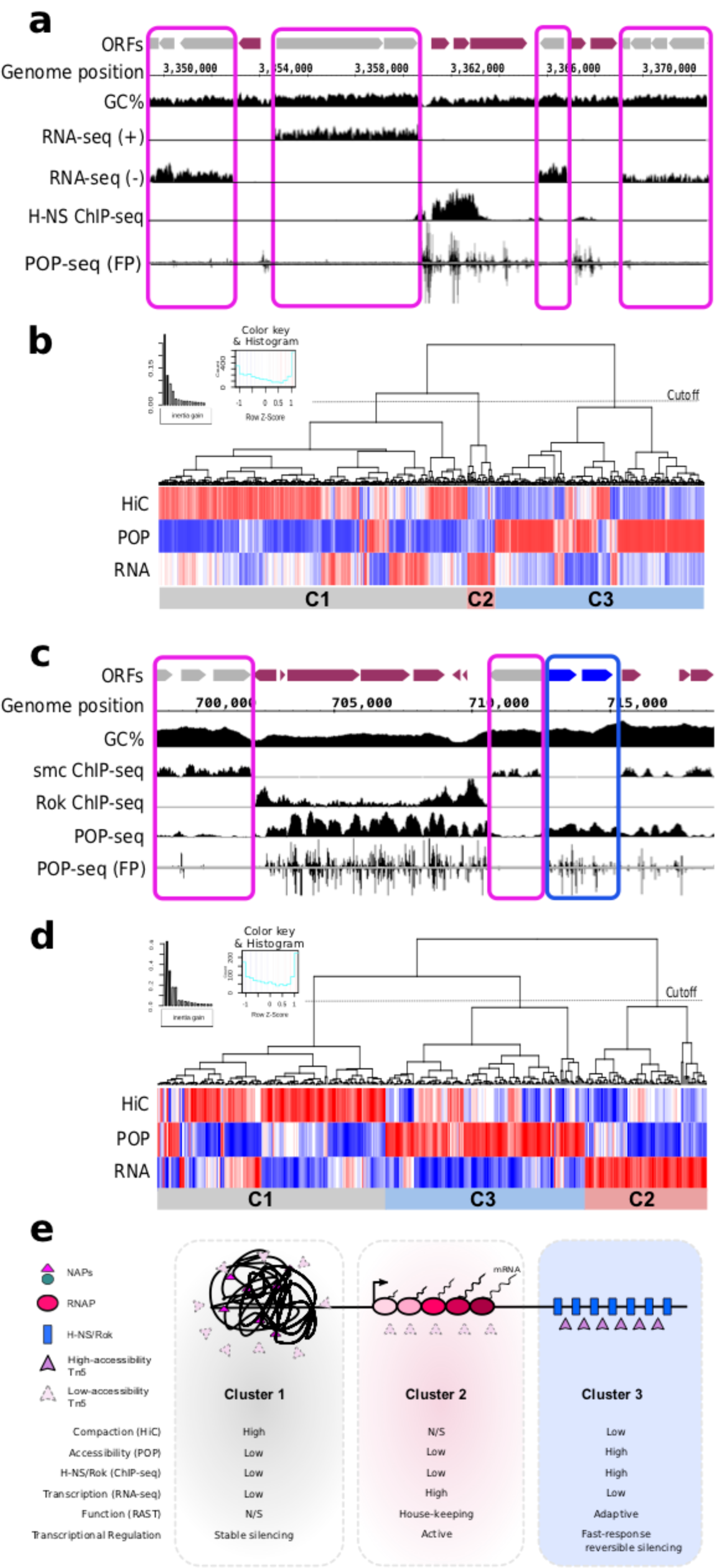
Relationship between transcription, nucleoid compaction, and Tn5 accessibility. **(a)** Comparison between the GC content, RNA-seq, H-NS Chip-seq, and POP-seq signals over a section of the *E. coli* genome. Actively transcribed genes (grey arrows highlighted with magenta boxes) show low POP-seq signal. Untranscribed genes show high H-NS binding and high POP-seq signals. **(b)** Heatmap and hierarchical clustering agglomeration (HCA) (Ward agglomeration on Euclidean distances) of 5 kb bins of the *E. coli* genome according to HiC, RNA-seq, and POP-seq signals. The three genomic clusters, C1, C2, and C3, are significantly enriched in HiC, RNA, or POP-seq signals, respectively. **(c)** Comparison between GC content, RNA-seq, Smc, and Rok Chip-seq and POP-seq signals over a section of the *B. subtilis* genome. Genes with high AT content show high Rok Chip-seq and POP-seq signals. SMC occupied regions are not accessible by Tn5 (magenta boxes and grey genes). Some regions are hypersensitive to Tn5 but the DNA binding protein is unknown (blue boxes and blue genes). (**d**) Heatmap and HCA of the 10 kb bins of the *B. subtilis* genome. Similar to *E. coli* (b), C1, C2, and C3 clusters are found, with significantly enriched HiC, RNA, or POP-seq signals, respectively. (**e**) Schematic view of the H-NS / Rok epigenetic control of the *E. coli* / *B. subtilis* genome highlighted by POP-seq experiment. Cluster 1 (C1) is characterized by a high genome compaction making it not accessible to neither Tn5 (low POP) nor RNA polymerase (low RNA-seq). Cluster 2 (C2) refers to regions with active transcription, contains mostly house-keeping genes, and the RNA-polymerase activity hinders Tn5 accessibility (low POP). Cluster 3 (C3) demonstrates a high POP and H-NS/Rok binding, which hinders RNA polymerase transcription. The H-NS regulated genes are involved in fast-responsive functions and depend strongly of the growth conditions.

Next, we performed a hierarchical cluster analysis (HCA) of the 928 *E. coli* genomic bins according to the POP-seq, RNA-seq, and HiC signals. Three distinct clusters (C1-3) were identified (**Fig. 4b**). C1 (56% of the *E. coli* genome) represents genomic regions with low openness, high compaction and low transcription, C2 (5% of the genome) represents highly transcribed regions, and C3 (38% of the genome) represents regions with high accessibility, low compaction and low transcription (p-value of Student’s t-test<0.05).

Our data show that the organization of the genomic regions can be driven either by compaction (C1), transcription (C2), or openness (C3), in a mutually exclusive fashion (**Fig. 4b**). We show that transcriptionally active regions (C2) are enriched in housekeeping functions (RAST analysis, Fisher test p-value<0.05). The high correlation between POP-seq and H-NS ChIP-seq data suggests that the gene expression in the accessible regions is silenced by H-NS. The C3 regions are significantly associated with alternative sugar utilization pathways (Fisher test p-value<0.05). The compacted regions (C1) had no significant enrichment of any biological function. Our results reinforce the notion that the genome structure-function relationship is governed by compaction, transcription, and openness, and the latter appears to include highly dynamic and fast-responsive functions.

### *POP-seq experiment of* B. subtilis *captures Rok transcriptional control*

We conducted POP-seq experiments in two biological replicates for *B. subtilis* to test the applicability of POP-seq across other bacterial phyla. The two POP-seq biological replicates were highly correlated (Pearson’s R=0.997, p-value<2.2e^-16^, 10-kb bins), confirming that, like *E. coli*, NAP and TF binding in *B. subtilis* is strictly regulated. We also observed that, as in *E. coli*, genes exhibiting the highest POP-seq signal are involved in fast-adaptive RAST functions (sporulation, sugar and amino acid utilization, multidrug resistance proteins), and the majority of less accessible genes were involved in housekeeping functions (translation initiation, cytochrome C and D, ribosomal proteins) (p-value of Mann-Whitney test, FDR adjusted<0.05).

Despite having comparable GC content (50.8% for *E. coli* and 43.6 % for *B. subtilis*), the genome organization of *E. coli* and *B. subtilis* is remarkably different. Rok and SMC (structural maintenance of chromosomes) play key roles in genome organization and are both among the best characterized NAPs in *B. subtilis*. Rok is a functional analogue of H-NS and ChIP-seq experiments have shown that Rok binds to AT-rich regions (Smits & Grossman 2010, **Fig. 4c**). The correlation between POP-seq and Rok ChIP-seq data was strongly significant (p-value<2.2^-16^). However, unlike in *E. coli* where H-NS and POP-seq signals were highly correlated (**Fig. 4c**), the Rok ChIP-seq signals were only partially correlated with the POP-seq signals (Pearson’s R=0.44, 10-kb bins).

The SMC complex is composed of Smc, ScpA and ScpB and is known to have a critical role in long-range DNA compaction^25^. Therefore, we explored the contribution of SMC binding to the total POP-seq signal by comparing our POP-seq data to previously published SMC ChIP-seq data^25^. We found that the POP-seq (and the majority of Rok ChIP-seq) signals are mutually exclusive to the SMC ChIP-seq signals (**Fig. 4c**), implying that SMC occupied regions are inaccessible to Tn5. Regions highly accessible to Tn5 that are neither occupied by Rok nor by SMC (**Blue box Fig. 4c**) raise the possibility of a hitherto unknown NAP that has binding properties reminiscent of Rok.

### B. subtilis *and* E. coli *have three major openness clusters in common*

To further investigate the relationships between genome accessibility, compaction, and transcription, we performed HCA for 10-kb bins of the *B. subtilis* genome for the POP-seq, HiC, and RNA-seq datasets. Similar to *E. coli*, our results led to the identification of three major clusters C1, C2, and C3, characterized by high HiC, high RNA, or high POP-seq signals, respectively (p-value of pairwise Student’s t tests, FDR adjusted<0.05) (**Fig. 4d**), confirming that Tn5 accessibility is constrained by compaction or transcription in *B. subtilis* as well as in *E. coli*.

Together, the results of HCA in *B. subtilis* show remarkable concordance with those obtained from *E. coli* (**Fig. 4d**), demonstrating the potential of POP-seq to study the open nucleoid of a wide range of bacterial phyla. Moreover, our observations in both bacteria also suggest that POP-seq can reveal the epigenetic mechanisms controlling the transcription of AT-rich genes that are required for fast environmental and nutritional adaptations (**Fig. 4e**). Indeed, HCA identified three different genomic clusters driven either by compaction (C1), transcription (C2), or openness to NAP binding/silencing (C3) in a mutually exclusive manner.

We suggest that in C1, the high level of compaction prevents both Tn5 accessibility and NAPs DNA binding, and occludes RNA transcription, suggesting a stable, condition-independent silencing of genes in those regions. Regions with active transcription (C2) are not accessible to Tn5 and are significantly enriched in housekeeping genes. Finally, highly accessible regions (C3) show a high AT content and H-NS/Rok binding, causing decreased transcriptional activity despite low compaction. These NAP-silenced genes are involved in fast-response functions and their expression level depends strongly on the growth condition, as shown by the differences between the POP profiles of *E. coli* grown in different culture media (**Fig. 3**).

## DISCUSSION

DNA folding proteins participate in genome structure organization, thus influencing DNA compaction, transcription, and replication^1^. The *in vivo* monitoring of these proteins on a genome-scale has so far been hindered by the lack of high throughput tools and the few tools currently available are laborious and limited to a handful of selected organisms. To address this technical gap, we developed POP-seq, which employs genome-wide Tn5 tagmentation to identify thousands of protein-DNA binding events *in vivo*. Integrating POP-seq with HiC and RNA-seq data showed that the *E. coli* and *B. subtilis* genomes are broadly organized into regions that are either compacted, highly transcribed, or open to the AT-binding NAPs (H-NS and Rok). Moreover, the protein-DNA binding events detected by POP-seq can be used to determine TFBS as well as for monitoring the genome openness as a proxy for overall nucleoid structural changes. Thus, we argue that POP-seq provides an essential new perspective on the bacterial nucleoid that could lead to in-depth understanding of the interplay between genome structure, specific functions and the overall phenotype.

Our results show a strong concordance between POP-seq signal and binding of important regulatory NAPs, as shown by the correlations between POP-seq, and H-NS in *E. coli*. In *B. subtilis*, the binding of Rok explains a major proportion of the observed POP-seq signal, but there are many AT-rich regions, which are accessible by Tn5, but are unoccupied by Rok and it is unclear whether there are other NAPs that bind to these regions. The genomic regions occupied by SMC are not accessible to Tn5 and DNA binding of both Rok and SMC seems to be mutually exclusive, hinting towards a novel functional role for Rok (and likely another unknown NAP) in constraining the genome-wide SMC binding regions.

POP-seq quantifies the concerted effects of all NAPs and TFs in the cell growing in a given environment with high sensitivity. By comparing these signals at different growth conditions, we can readily detect changes in the nucleoid structure. Thus, we propose that POP-seq constitutes an accurate, high throughput and cost-effective method for the study of protein-DNA interactions. POP-seq can be applied to identify active TF and NAP binding sites *in vivo* with high accuracy, thereby allowing a system-level understanding of the dynamics underlying gene regulatory networks.

Functional analysis of the most- and least-accessible genes shows that the nucleoid structure is carefully controlled to achieve an optimal transcriptional profile. Moreover, multi-omics integration of genome-wide POP-seq, ChIP-seq, HiC, and RNA-seq data provided insights regarding the interplay between genome accessibility, DNA compaction, and RNA transcription in control of gene expression and distinguishing phenotypes. This view is supported by the finding that *E. coli* genome accessibility is greatly modified after growth in the presence or absence of glucose as carbon source, with important decrease of POP-seq signal (likely due to the removal of H-NS binding) upon activation of alternative sugar utilization pathways. We thus demonstrate that POP-seq aids in unraveling epigenetic control of genes requiring a fast and reversible adaptation to the environment, by the competition between RNA polymerase and NAPs for DNA binding.

In summary, POP-seq enables rapid elucidation of the openness of the prokaryotic chromatin, which is directly linked to its structure. POP-seq is independent of culture synchronization and provides high resolution mapping of protein-DNA events. Due to its simplicity and cost effectiveness, it can be implemented to study a plethora of bacterial species in high throughput to elucidate the structural changes in the chromatin and link them directly to phenotype.

## METHODS

### Bacterial strains and growth conditions

*E. coli* K12 substrain MG1655 was grown in LB for most of the experiments. Other *E. coli* growth media used in this study is M9 MM supplemented with either glucose, xylose or ribose. *B. subtilis* PY79 was grown in rich CH medium^26^. All bacterial cultures were done at 37°C with shaking and harvested at mid-log phase.

### POP-seq Method

Bacterial cultures were grown to mid exponential phase (OD600 = 0.3-0.5) in Luria-Bertani medium at 37°C. Crosslinking was achieved by treatment with 1% formaldehyde for 20 minutes. Cells were pelleted by centrifugation and the cell pellets were lysed by grinding in liquid nitrogen or by bead bashing at 4°C for 10 mins. 500 µL SET buffer (75 mM NaCl, 25 mM EDTA pH 8, 20 nM Tris-HCl pH 7.5) were used for grinding. Lysate was resuspended in 2x protease inhibitor solution (cOmplete mini, Roche) and centrifuged for 10 min at 14,000 rpm and 4°C. 25 µL of supernatant was used for buffer exchange with Tris-EDTA (10 mM Tris, 1 mM EDTA, pH 8) with a 45 min incubation period at room temperature.

Hi-sensitivity Qubit DNA kit (Thermo Fisher) was used to measure the DNA concentration directly from the lysate. 700 pg DNA were used as input for the illumina Nextera kit. After library preparation, AMPure beads were used to purify the library as recommended by the manufacturer. The libraries from all experiments were sequenced in either the Illumina HiSeq™ 4000 or MiSeq™ instruments at UCSD IGM genomics center. 100 bp-cycle kits or 150 bp-cycle kits were used to sequence the libraries.

### Data preparation for POP-seq versus HiC comparison

The *E. coli* HiC contact matrices (GSM2870426_mat_BC110_CACT_wt_LB_37C.txt.gz and GSM2870427_mat_BC110_CACT_wt_LB_37C_rep1.txt) were both acquired from Lioy *et al*. (2018)^11^. The mean one-dimensional (1-D) structure of the *E. coli* genome was reconstructed from the 3-D contact data, GitHub repository https://github.com/koszullab/E_coli_analysis/blob/master/python_codes/comp_short.py^11^ accessed Nov. 13^th^ 2019). The resulting 1-D HiC data had a total 928 bins. However, an intrinsic curvature, which hinders comparison with other datasets was seen. Accordingly, we corrected the HiC data by local regression fitting model (**Supplementary Fig. 1**) on R version 3.4.4 (R Core Team, 2018). Each bin had RPM values summed over windows of 5000 nucleotides. The resulting dataset was z-score-normalized and smoothed using the Signal.savgol_filter from the Scipy Python library with window_length of 11 for POP-seq and 7 for HiC (due to its lower resolution) and a polyorder of 3.

The 2D matrices of the *B. subtilis* HiC contact maps (GSE68418, GSM1671399_01_Rudnerlab_HindIII_HiC_PY79.matrix.txt.gz and GSM1671400_02_Rudnerlab_HindIII_HiC_PY79_rep1.matrix.txt.gz) were acquired from Wang et al. (2015)^10^. The 3D mean matrix was flattened to a 1D array as in *E. coli*, without the need to correct for curvature. However, the resulting bins were only 404 bins, due to the 10-kb resolution of the original HiC dataset. The resulting dataset was z-score-normalized and smoothed using the Signal.savgol_filter from the Scipy Python library with window_length of 15 for POP-seq and 25 for HiC (due to its lower resolution) and a polyorder of 3.

### ChIP-seq and RNA-seq and data analysis and plotting

The H-NS ChIP-seq sequencing files (ERR01957) were acquired from Kahramanoglou et al. (2011)^18^. The *B. subtilis* Rok ChIP-seq was acquired from Smits and Grossman (2010)^6^ (GSE23199). The *B. subtilis* SMC ChIP-seq was acquired from Wang et al. (2017)^25^ (GSE85612). The *E. coli* RNA-seq data was acquired from Choi et al. (2019)^24^ (SRR8242101 and SRR8242105). The *B. subtilis* RNA-seq data was acquired from SRR2984942-3 under accession number PRJNA304431. Bowtie2 was used to align the sequencing reads using default parameters. The coverage was calculated with samtools mpileup and normalized in RPM (reads per million) and the resulting wig files were plotted using Integrated Genome Browser (IGB).

### Data analysis

In the general procedure, primers and adapter sequences were removed using trim_galore (https://www.bioinformatics.babraham.ac.uk/projects/trim_galore/) in paired-end mode (--paired) with the quality cutoff (-q) set to 22 and -fastqc enabled. Next, reads were aligned to the reference genome using bowtie2^27^. Wig files containing the number of mappings at each genome position were generated using the samtools mpileup command and normalized by reads per million (RPM). The resulting wig files were processed using in-house Python scripts. FeatureCounts^28^ was used to determine the number of fragments corresponding to each region of interest (features), which could be a gene or promoter. A minimum of 2/3 of each read must be within the gene in order for it to be assigned (--fracoverlap 0.66). DESeq2^29^ was then used to determine the differential POP-seq and RNA-seq signals for each feature. Downstream analyses were performed using R version 3.6.1 (R Core Team, 2019).

### POP-seq footprinting

In order to accurately determine the location of specific Tn5 transposition events so as to precisely pinpoint individual binding sites, mapped reads were trimmed to a 9 bases at the 5′ end which are normally covered by the Tn5 protein at the site of transposition^13^. Trimming and alignment were performed as in the general case using trim_galore and bowtie2.

### Statistical analyses

Statistical analyses were performed using R version 3.6.1 (R Core Team, 2019). All statistical tests were considered significant if the p-value was below 0.05, after False Discovery Rate (FDR) adjustment in case of multiple testing^30^. A logistic regression supervised model was built to predict the likelihood of each nucleotide to be at the vicinity of a TFBS according to POP-seq measurements. For each nucleotide, the dependent variable was coded 1 if the corresponding genome position was referenced as a TFBS in EcoCyc^17^, 0 if not. Explanatory variables were selected by stepwise bottom-up selection according to the Akaike Information Criterion^31^, and included POP-seq and H-NS ChIP-seq signals, as well as the presence/absence of a gene in every genome position. The logistic regression returns a value in the interval [0, 1] and the best threshold for setting the prediction to 1 (event) or 0 (no event) for each genome position was calculated by analysis of the receiver operating characteristic (ROC) curve. The logistic model was built using the R package MASS and ROC curve analysis was performed using the pROc package.

## ACKNOWLEDGEMENTS

This study is based on work supported by the U.S. Department of Energy (DOE), Office of Science, Office of Biological & Environmental Research under Awards DE-SC0012658 and DE-SC0012586.

## AUTHOR CONTRIBUTIONS

M.M.A-B and K.Z. conceived the study with input from O.M. M.M.A-B, N.C. O.M. and N.C. carried out the experiments and performed all data analysis. All authors wrote and reviewed the manuscript.

## SUPPLEMENTARY DATA

**Supplementary figure 1.**
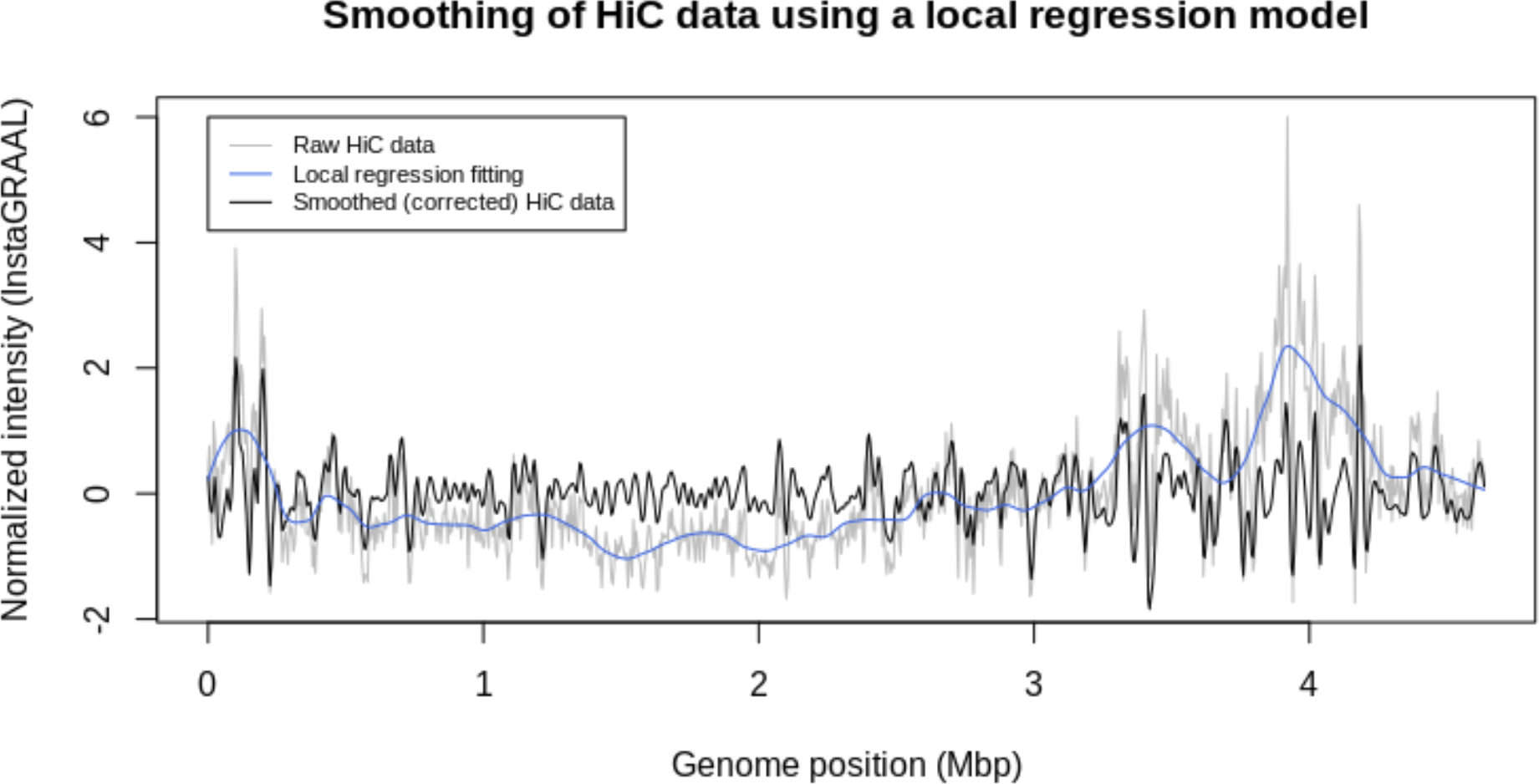
Smoothing of HiC data using a local regression fitting. We used a local regression model (blue line) of the raw HiC data (grey line) to calculate smoothed HiC values (black line).

**Supplementary table 1.** Top 20 subsystems with the most significantly higher POP signal.

| *Subsystem* | *mean signal in subsystem* | *mean signal out of subsystem* | *p-value* | *p-value (adjusted)* |
| --- | --- | --- | --- | --- |
| LOS core | 26.01 | 4.40 | 0.00 | 0.00 |
| oligosaccharid |  |  |  |  |
| e biosynthesis |  |  |  |  |
| The usher | 31.08 | 4.46 | 0.00 | 0.00 |
| protein HtrE |  |  |  |  |
| fimbrial |  |  |  |  |
| cluster |  |  |  |  |
| CRISPRs | 16.38 | 4.52 | 0.00 | 0.01 |
| Mediator of | 9.30 | 4.54 | 0.00 | 0.02 |
| hyperadheren |  |  |  |  |
| ce YidE in |  |  |  |  |
| Enterobacteria |  |  |  |  |
| and its |  |  |  |  |
| conserved |  |  |  |  |
| region |  |  |  |  |
| Orphan | 7.54 | 4.53 | 0.00 | 0.02 |
| regulatory |  |  |  |  |
| proteins |  |  |  |  |
| Periplasmic | 52.86 | 4.46 | 0.00 | 0.03 |
| Acid Stress |  |  |  |  |
| Response in |  |  |  |  |
| Enterobacteria |  |  |  |  |
| D-gluconate | 9.03 | 4.53 | 0.00 | 0.03 |
| and |  |  |  |  |
| ketogluconate |  |  |  |  |
| s metabolism |  |  |  |  |
| D-allose | 11.44 | 4.53 | 0.00 | 0.05 |
| utilization |  |  |  |  |
| General | 6.40 | 4.55 | 0.00 | 0.05 |
| Secretion |  |  |  |  |
| Pathway |  |  |  |  |
| The fimbrial<br>Sfm cluster | 23.00 | 4.52 | 0.00 | 0.05 |
| Xylose<br>utilization | 11.12 | 4.53 | 0.00 | 0.06 |
| Biofilm<br>Adhesin<br>Biosynthesis | 10.72 | 4.54 | 0.00 | 0.06 |
| Curli<br>production | 12.71 | 4.53 | 0.00 | 0.07 |
| L-fucose<br>utilization | 7.18 | 4.55 | 0.00 | 0.07 |
| A toxin-antitoxin<br>module<br>cotranscribed<br>with DinB | 23.88 | 4.53 | 0.00 | 0.07 |
| Lysine<br>degradation | 18.18 | 4.53 | 0.00 | 0.07 |
| D-ribose<br>utilization | 8.74 | 4.54 | 0.00 | 0.07 |
| The fimbrial<br>Stf cluster | 9.34 | 4.54 | 0.01 | 0.08 |
| Unknown<br>carbohydrate<br>utilization<br>(cluster Ydj) | 7.52 | 4.54 | 0.01 | 0.09 |
| L-rhamnose<br>utilization | 8.12 | 4.54 | 0.01 | 0.09 |

**Supplementary table 2.** Top 20 subsystems with the most significantly lower POP signal.

| *Subsystem* | *mean signal in subsystem* | *mean signal out of subsystem* | *p-value* | *p-value (adjusted)* |
| --- | --- | --- | --- | --- |
| Ribosome<br>LSU bacterial | 1.85 | 4.59 | 0.00 | 0.00 |
| Ribosome<br>SSU bacterial | 0.84 | 4.59 | 0.00 | 0.00 |
| Respiratory<br>Complex I | 0.89 | 4.58 | 0.00 | 0.00 |
| Na(+)-<br>translocating<br>NADH-quinone<br>oxidoreductas<br>e and rnf-like<br>group of<br>electron<br>transport<br>complexes | 0.97 | 4.57 | 0.00 | 0.01 |
| Alkylphospho<br>nate<br>utilization | 1.96 | 4.57 | 0.00 | 0.06 |
| Ethanolamine | 2.23 | 4.57 | 0.00 | 0.06 |
| utilization |  |  |  |  |
| Mycobacteriu | 0.28 | 4.56 | 0.00 | 0.06 |
| m virulence |  |  |  |  |
| operon |  |  |  |  |
| involved in |  |  |  |  |
| protein |  |  |  |  |
| synthesis |  |  |  |  |
| (LSU |  |  |  |  |
| ribosomal |  |  |  |  |
| proteins) |  |  |  |  |
| Histidine | 1.38 | 4.57 | 0.00 | 0.06 |
| Biosynthesis |  |  |  |  |
| Thiamin | 1.79 | 4.57 | 0.00 | 0.06 |
| biosynthesis |  |  |  |  |
| Osmoprotecta | 1.29 | 4.56 | 0.01 | 0.09 |
| nt ABC |  |  |  |  |
| transporter |  |  |  |  |
| YehZYXW of |  |  |  |  |
| Enterobacteria |  |  |  |  |
| les |  |  |  |  |
| De Novo | 1.80 | 4.57 | 0.01 | 0.13 |
| Purine |  |  |  |  |
| Biosynthesis |  |  |  |  |
| Fatty Acid | 2.22 | 4.57 | 0.02 | 0.17 |
| Biosynthesis |  |  |  |  |
| FASII |  |  |  |  |
| tRNA | 0.40 | 4.56 | 0.02 | 0.17 |
| aminoacylation, Phe |  |  |  |  |
| Transcription initiation, bacterial sigma factors | 1.85 | 4.57 | 0.02 | 0.20 |
| Bacterial Cytoskeleton | 2.06 | 4.57 | 0.03 | 0.22 |
| The mdtABCD multidrug resistance cluster | 1.46 | 4.56 | 0.03 | 0.22 |
| KDO2-Lipid A biosynthesis | 2.34 | 4.57 | 0.03 | 0.22 |
| TCA Cycle | 1.87 | 4.56 | 0.03 | 0.22 |
| Biogenesis of c-type cytochromes | 2.65 | 4.56 | 0.03 | 0.22 |
| RuvABC plus a hypothetical | 1.22 | 4.56 | 0.03 | 0.24 |

